# Efficient detection of pre-proinsulin by double antibody sandwich ELISA

**DOI:** 10.1101/2020.03.27.011098

**Authors:** Zhu Zhu, Guoliang Ma, Li Wang, Han Wang, Hengfang Tang, Zhou Wei, Peng Wang

## Abstract

To generate monoclonal antibodies against pre-proinsulin (PPI), and establish sandwich ELISA method to provide a basis for PPI detection in recombinant human insulin production.

**Methods:** The Balb /c mice were immunized with PPI, and the hybridomas secreting anti-PPI monoclonal antibodies were obtained by conventional cell fusion technique and ELISA screening.The antibody was purified using a Protein G gel column and identified for purity by SDS-PAGE. Pairing effect was found by the sandwich ELISA, and the specificity of the paired antibody was determined. A paired antibody with better specificity was selected to establish sandwich ELISA, and construct a quantitative curve, the accuracy and sensitivity of the method were evaluated.

**Results:** Six anti-PPI monoclonal antibodies were obtained, named P1, P2, P3, P4, P5 and P6, of which P5 had the highest titer. The sandwich ELISA method was established with P5 for plating and P2 for detection antibodie. The linear range of the quantitative curve of PPI by sandwich ELISA was 0. 645-82.5 pg/mL, the recovery was 89%–95%, and the limit of detection was 3.06 pg/mL.

**Conclusion:** Six monoclonal antibodies against PPI were generated and the sandwich ELISA method was established to detect PPI in process control and product release control for recombinant human insulin production.

---

Insulin regulates glucose metabolism and controls blood sugar balance are the main drugs used in the treatment of diabetes^[1]^. At present, high-purity insulin is obtained by enzymatic digestion and purification of inactive pro-proinsulin (PPI)^[2–3]^. The efficient and stable expression of PPI is a prerequisite for obtaining high yield insulin. However, PPI that has not been digested and removed will seriously affect the quality and safety of insulin^[4–5]^.Therefore, it is very important to accurately detect the expression of PPI and the residual amount of PPI in insulin.

The European Pharmacopoeia clearly requires that PPI in insulin must be detected and controlled, but no specific detection method has been proposed. At present, the commonly used methods are radioimmunoassay, high performance liquid chromatography and so on^[6–8]^. Radioimmunoassay methods have been gradually phased out due to inadequate prices, prone to environmental pollution, and harm to human health^[9]^. High performance liquid chromatography is often used to detect the PPI content in the purification process, but the sensitivity is poor and it cannot meet the requirements for the detection of PPI residues in the final product^[10]^.

Enzyme-linked immunosorbent assay has high sensitivity, which can be used in the detection of trace PPI. However, due to the large differences in PPI produced by different processes, it is currently impossible to use a universal ELISA kit for determination. In this study, the PPI molecule consists of A-chain of insulin, B-chain of insulin, the guide peptide (SOD) and the lysine and arginine (-Lys-Arg-) linker. After trypsin digestion, SOD and single chain Lys-Arg-insulin (single chain Lys-Arg-Insulin, sIKR) were produced. As the enzyme digestion proceeded, the junction of A chain and arginine was cut open, only the remaining B chain was linked to Arg-Lys-, and the free A chain was correctly paired with B chain to form Arg-Lys-insulin (IKR). ikr formed lys-insulin (ik) after carboxypeptidase b removed the arginine residue, and ik further removed the lysine residue to form insulin with the correct spatial conformation. To find a highly targeted and sensitive method, six anti-pi monoclonal antibodies were prepared by hybridoma technique, and a double antibody sandwich elisa was preliminarily established to determine the ppi residue in recombinant human insulin.

## 1. Materials and Methods

### 1.1 Materials

Preparation, purification and labeling of PPI monoclonal antibody: experimental animals were 6 to 8 weeks old Balb/c mice, penicillin, streptomycin and fetal bovine serum were purchased from Gibco, and horseradish peroxidase labeled sheep anti-rat IgG (HRP-sheep anti-mouse IgG) was purchased from Ferdbio, HAT (50×), HT, PEG1450 and Incomplete freund’s adjuvant Purchased from Sigma, DMEM Medium purchased from Wisent, cell dissociation buffer (0. 25% pancreatin and 0. 02%EDTA) preparation for use. The reagents used for antibody purification including antibody elution buffer, neutralization buffer, preservation buffer, pbs (ph 7.4) buffer, and sodium azide were self-made.

The reagents used for elisa detection in the establishment of the double antibody sandwich method are as follows: elisa coating solution, washing solution (pbst),3,3’,5’-tetramethylbenzidine (3,3’,5’-tetramethylbenzidine, hereinafter referred to as tmb) substrate reaction solution, sealing solution, universal dilution solution, termination solution are all self-made.

Animal immune antigen ppi, indirectly elisa method coated antigen ppi, double antibody sandwich method ppi standard and sod, sikr, ikr, ik are collected recombinant human insulin production process ppi and its digested intermediate product after purification, the purity is over 95%.

### 1.2 Methods

#### 1.2.1 Preparation, purification and marking of monoclonal antibodies

PPI was used as immunogen immunized 6- to 8-week-old balb/c mice and blood was taken before immunization as a negative serum control. the initial immunization dose was 100μg/ only, and the mice were immunized after emulsification with the same volume of ferrin complete adjuvant. after every 2 weeks, the serum antibody titer was detected two weeks after the third immunization. the mice with higher titer were immunized by intraperitoneal injection of immunogen 100μg/ only 3 days before cell fusion. Then the blood was sacrificed and the immune spleen cells were isolated. The immune spleen cells and the already prepared mouse myeloma cells sp2/0 were fused. ELISA screen positive monoclonal hybridoma cells, after three subclonal and ELISA test completely positive, expand the culture and build the plant. The screened hybridoma cells with stable secreted target antibody were inoculated with 1 to 5×106/ only in balb/c mice aged 6 to 8 weeks with incomplete frst adjuvant. When the abdomen of the mice was obviously enlarged and the skin was tense at the touch of the hand, the ascites were collected with 10 ml needle. Centrifugation of ascites, collection of supernatant, determination of titer. purified with ProteinG gel column and purified monoclonal antibody was identified by SDS-PAGE gel electrophoresis. The obtained monoclonal antibody was diluted to 1 mg/mL with 0.1 M PBS buffer (pH 7.4), and the interfering substance was removed by ultrafiltration, adding water-soluble biotin, and reacting at room temperature for 1 hour. The free biotin was removed by ultrafiltration.

#### 1.2.2 Double antibody sandwich Elisa screening for paired antibodies

The purified mabs were diluted to 1 ug/ml with pbs (ph 7.4) and 100 ul/wells were separately encapsulated in the enzyme labeled plates. 2% BSA,150 ul/pore, closed at 37°C for 60 min, TBST washed 3 times. PPI with 1 ug/ml in the detection hole,100 ul/ pore, with 1% bsa as the blank control, was placed at 37°c for 60 min and washed by tbst for 3 times. arranged according to the checkerboard method, each monoclonal antibody labeled with biotin was added,100 ul/pore, placed at 37°c for 60 min, and tbst was washed 3 times. plus streptavidin hrp,100 ul/pore, placed at 37°c for 60 min. Tbst washing 3 times, adding tmb color rendering solution to avoid light for 5 min, adding terminal solution to read the absorbance value at 450 nm wavelength. The sample hole absorbance value is more than 2.1 times the blank control hole absorbance value as the positive detection hole.

#### 1.2.3 Specific evaluation of paired antibody

The coating antibody corresponding to the positive detection hole in 1.2.2 and the biotin-labeled antibody were used as the solid-phase antibody and the detection antibody, respectively. ppi, sod, sikr, ikr and ik of 1ug/ml were added in the detection hole, and 1% bsa was used as the blank control to test the specificity of each pair antibody according to the method described in 1.2.2.

#### 1.2.4 Construction of quantitative curve

The determined optimum solid-phase antibodies were coated in the enzyme labeled plate with 100 μ L series concentrations of PPI:82.5 PG/mL,41.25 PG/mL,20.65 PG/mL, 10.31 PG/mL,5.16 PG/mL,2.58 PG/mL,1.29 PG/mL,0.645 PG/mL,0 PG/mL,60 min at 37 ° C, and TBST washing 3 times. then add 1ug/ml,100ul/pore optimum detection antibody, follow the method described in 1.2.2 for the operation of double antibody sandwich elisa with 3 multiple holes per ppi concentration. With the average od450 of each concentration as the ordinate and the ppi concentration as the abscissa, the standard curve was fitted with skan itre to determine the linear range and quantitative equation of the detection.

#### 1.2.5 Accuracy and sensitivity evaluation

The recombinant human insulin was accurately weighed, and 2ug/ml was prepared with universal diluent, and 2ug/ml recombinant human insulin solution 200ul was added 82.5pg/mL,20.58pg/mL and 5.16pg/mL respectively. Dilute the 2ug/ml recombinant human insulin solution to 1ug/ml as a sample to be tested. the standard sample, the sample to be inspected, the quality control sample were added into the elisa board of the package to be completed, and the universal diluent was used as the blank, each sample 3 multiple holes, according to the above established elisa method. The accuracy is expressed by the added recovery rate, that is, the average concentration of the measured quality control sample minus the ratio of the difference of the average concentration of the measured sample to the theoretical added concentration (%).

The absorbance of 12 blank samples, the general dilution at 450 nm wavelength, was determined by the double antibody sandwich ELISA method, and the average value of the measured samples plus the absorbance obtained by 3 times standard deviation was substituted into the quantitative equation.

## 2. Results and Discussion

### 2.1 Preparation and purification of monoclonal antibody

Using ppi-immune mice, hybridoma cells were prepared. after multiple rounds of screening and three subclones, six cell lines that stably secreted anti-pi monoclonal antibodies were obtained, namely p1, p2, p3, p4, p5, and p6. in subsequent studies, each monoclonal antibody was named after its source cell line. the selected cell lines were injected into the mouse peritoneal cavity with a quantity of 1 5 106/only to prepare ascites, and the collected ascites supernatant was measured with titer (figure 1). he results showed that p5 had the highest titer, p1 titer and p3 titer were the lowest.

**Fig. 1.**
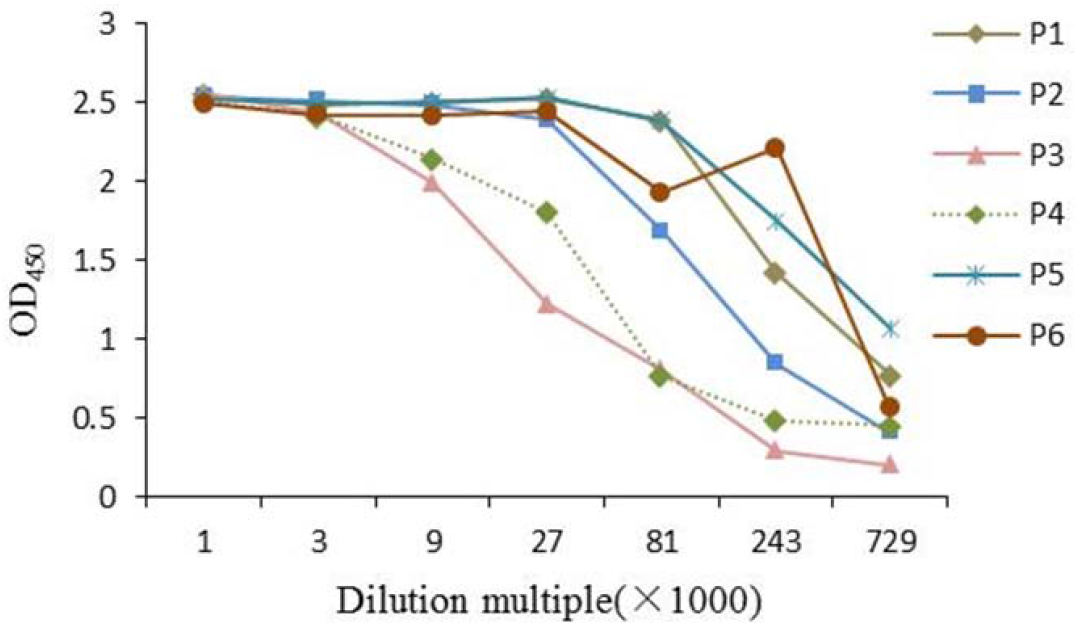
Titers of monoclonal antibodies against PPI

However, in general, each single titer is above 1:7.29×105, which can be further studied. Protein G column was used for affinity chromatography of monoclonal antibody P1, P2, P3, P4, P5, and P6, and purified monoclonal antibody was detected by reducing SDS-PAGE (Fig. 2). The results showed that each monoclonal antibody had high purity and no miscellaneous protein except immunoglobulin light chain and heavy chain.

**Fig. 2.**
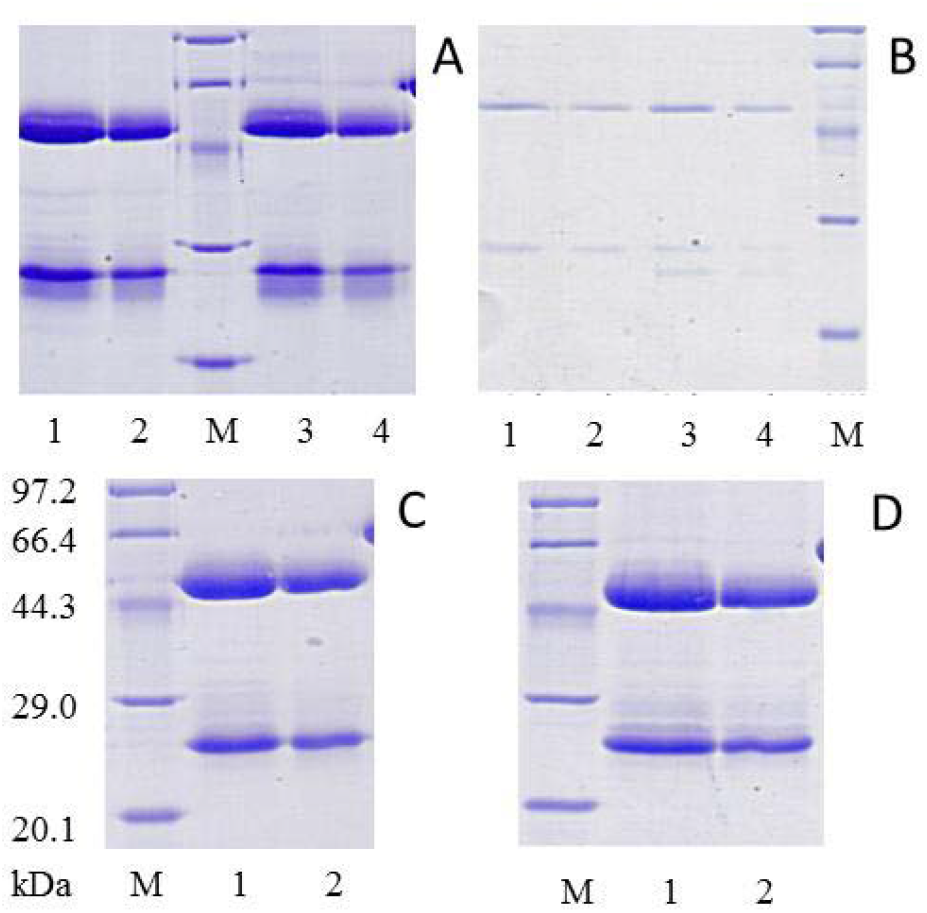
Purification of monoclonal antibodies against PPI. (A) Reduced SDS-PAGE showing purification of monoclonal antibodies P1 and P2. 1: 10 μL of P1; 2: 5 μL of P1; 3: 10 μL of P2; 4: 5 μL of P2;(B) P3 and P4. 1: 10 μL of P3; 2: 5 μL of P3; 3: 10 μL of P4; 4: 5 μL of P4; (C) P5. 1: 10 μL of P5; 2: 5 μL of P5; (D) P6. 1: 10 μL of P6; 2: 5 μL of P6; M: molecular weight mass standards (kDa).

### 2.2 Double antibody sandwich ELISA screening for paired antibodies

P1, P2, P3, P4, P5, and P6 were successively used as solid-phase antibody packages in the enzyme labeled plate, PPI was added as antigen and 1% BSA as blank control, and biotin-labeled monoclonal antibody was added for double antibody sandwich ELISA detection, respectively. the antibody corresponding to the detection hole with 2.1 times greater than the absorbance of the blank hole was used as the positive paired antibody (table 1). The results showed that when P5 was used as solid phase antibody, biotin-labeled P1, P2, P3, P4 and P6 could be used as detection antibody. P2 was not successfully paired as a solid antibody. P1, P3, P4 and P6 can only be detected with biotin-labeled P5 as solid-phase antibodies.

**Table 1.** Screening of paired antibodies

|  | P1 | P2 | P3 | P4 | P5 | P6 |
| --- | --- | --- | --- | --- | --- | --- |
| P1-bio | - | - | - | - | + | - |
| P2-bio | - | - | - | - | + | - |
| P3-bio | - | - | - | - | + | - |
| P4-bio | - | - | - | - | + | - |
| P5-bio | + | - | + | + | - | + |
| P6-bio | - | - | - | - | + | - |

### 2.3 Specific evaluation of paired antibody

P1, P3, P4 and P6 were used as solid-phase antibodies, and the intermediates SOD, sIKR, IKR, and IK were digested by PPI and P5 as antigens, respectively. The specificity of each paired antibody was determined (Fig. 3a). The results showed that when P3 was used as solid-phase antibody, the blank control absorbance was the highest, and the paired antibody did not react with each antigen; when P4 was used as solid-phase antibody, the paired antibody could react with PPI, sIKR and IKR, and the specificity was poor; P1 as solid-phase antibody, the paired antibody only reacted with PPI, but the absorptivity value was only 0.293; P6 as solid-phase antibody, the paired antibody produced immune response to both PPI and sIKR, and the absorbance value of sIKR was much higher than that of PPI 0.58. p5 was used as solid-phase antibody and ppi and its digested intermediates sod, ik, ikr and sikr were used as antigens. p1, p2, p3, p4 and p6 were used as detection antibodies to determine the specificity of each paired antibody (figure 3b). The results showed that the specificity of paired antibody was poor when P4 and P6 were used as detection antibody, and the absorbance value was close to that of PPI, sIKR, IKR and IK, respectively. When P3 was used as detection antibody, the highest absorbance value of IK was 2.47 and 2.46 respectively, but the positive value of paired antibody was 0.56,0.861 and 0.61 respectively, which was much lower than PPI, and the interference of PPI was smaller, so P5 antibody was used as solid phase antibody and P2 as detection antibody.

**Fig. 3.**
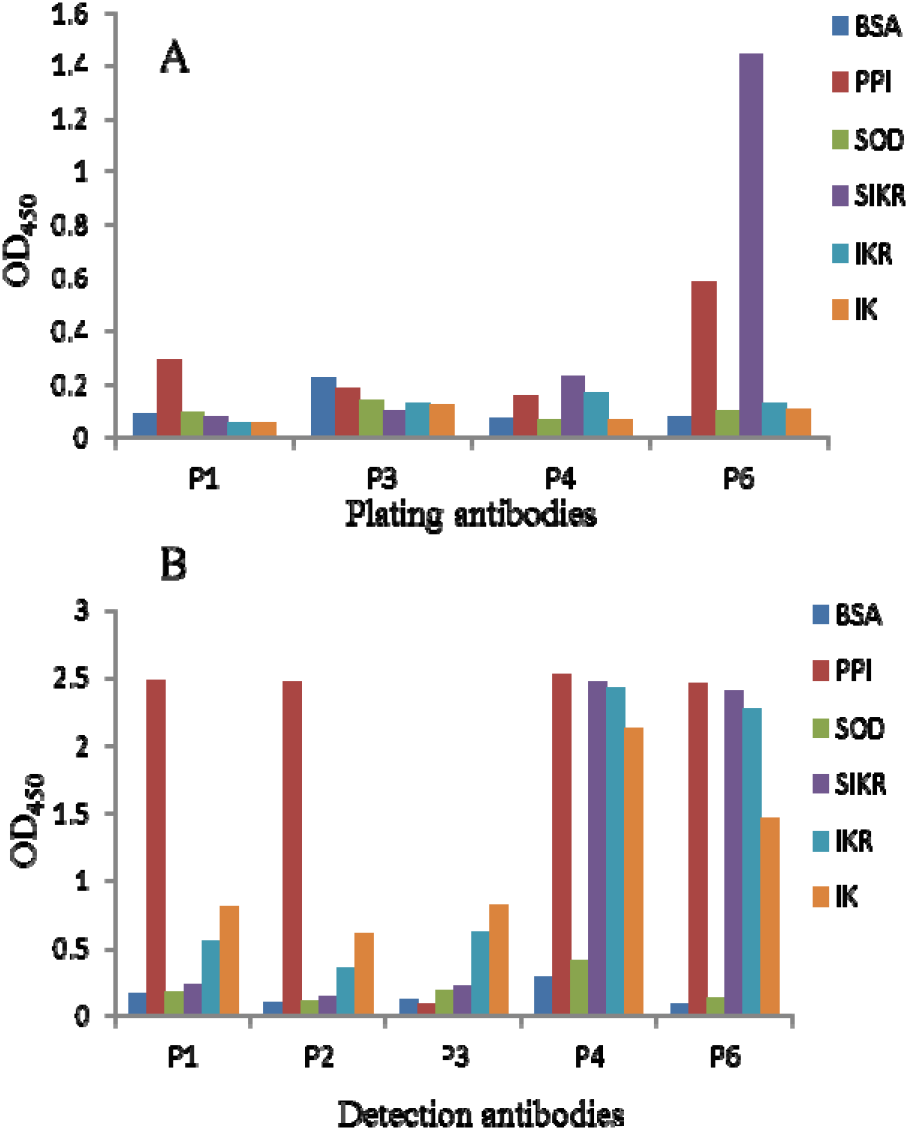
Specificity evaluation of paired antibody. (A) Plating antibodies are listed below the x-axis, detection antibodie is P5. (B) Plating antibodie is P5, detection antibodies are listed below the x-axis.

### 2.4 Construction of quantitative curves

P5 as a solid antibody, adding a series of concentration of PPI standard, using biotin-labeled P2 as the detection antibody, double antibody sandwich Elisa to determine PPI. Using OD450 as vertical coordinate and PPI concentration as horizontal coordinate, four-parameter fitting is carried out, and the standard curve is drawn (Fig. 4), and the fitting curve equation is obtained: y = 7.92579 + [(0.130957 – 7.92579) / (1 + (x/96.6272)^1.51852)] (R2=1), The detection method has a linear range of 0.645-82.5 PG/mL.

**Fig. 4.**
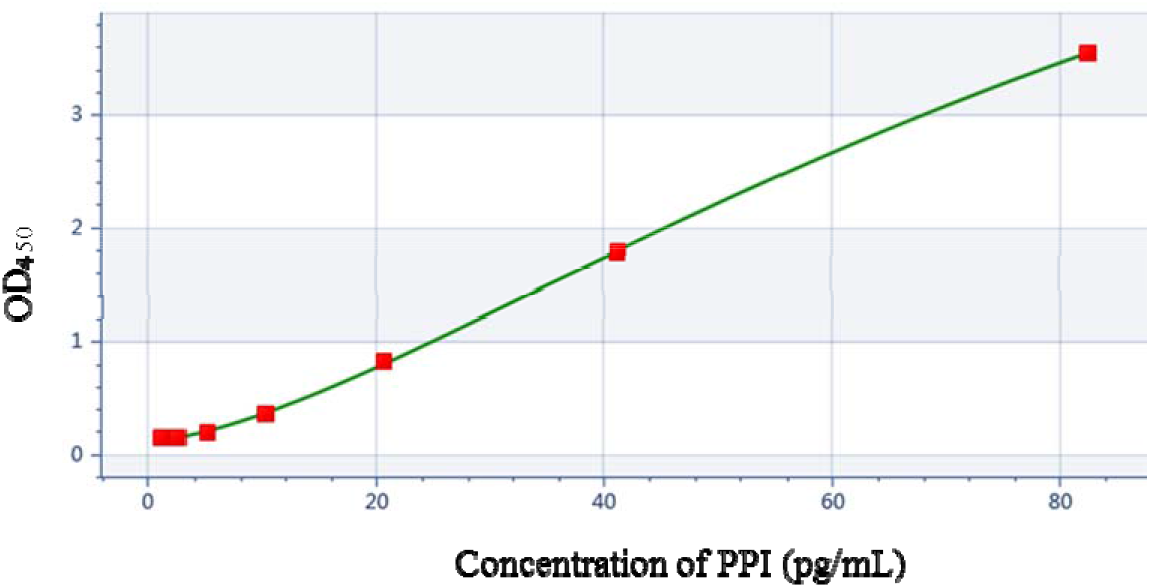
Quantitative curve of PPI

### 2.5 Accuracy and sensitivity evaluation of PPI method for double antibody sandwich ELISA

The recovery of quality control samples with high (41.25 pg/mL), medium (10.31 pg/mL) and low (2.58 pg/mL) concentrations were detected according to the accuracy determination method (Table 2). The results showed that the concentration of ppi measured in 1ug/ml recombinant human insulin solution was 0.638 pg/ml, and the recovery rate of each quality control sample was 91.4%,88.62% and 95.41%, respectively, with good accuracy.

**Table 2.** Recoveries of PPI from the samples fortified with different amounts of PPI

| Theoretical<br>concentration(pg/mL) | Measured<br>concentration(pg/mL) | Sample<br>concentration(pg/mL) | Recovery rate(%) |
| --- | --- | --- | --- |
| 41.25 | 38.34 |  | 91 |
| 10.31 | 9.78 | 0.638 | 89 |
| 2.58 | 3.10 |  | 95 |

According to the method of sensitivity determination, the OD450 values of 12 blank samples, that is, sample dilutions, were determined, the average value of the value obtained by adding 3 times standard deviation was 0.172, and the minimum detection limit was calculated to be 3.06pg/mL, which met the requirement of PPI detection sensitivity in recombinant human insulin.

## 3. Conclusion

In addition to the european pharmacopoeia 2020 edition of the chinese pharmacopoeia recombinant human insulin also proposed to control the production process of insulin precursors, it can be seen that the detection of insulin precursors has become an indispensable link in process control and final product release. The process control is mostly carried out by proper HPLC method^[11]^, but because of the low content of precursor material in the final product, the sensitivity of the detection method is high, which becomes the difficulty in the product release control detection of each enterprise.

The enzyme-linked immunosorbent assay was highly sensitive, but mostly used for the detection of natural insulin pro in serum^[12–13]^. In contrast, there are many kinds of intermediate products and similar molecular structure in the detection of precursor substances in insulin, which are prone to cross reactions, as well as the high concentration of insulin in the detection samples, and easy to interfere with the detection results. In this study, the purified insulin precursor was collected from the production process of recombinant human insulin in hefei tianmai biology as antigen to prepare monoclonal antibody. The monoclonal antibody was screened for paired antibody with two pairs, and the specific progressive detection of paired antibody was used to screen the paired antibody with better specificity. in addition, the accuracy as well as sensitivity of the method were verified. compared with the method established by leng et al., the sensitivity was significantly improved, and the insulin interference was excluded in the accuracy verification^[14]^, which initially established a more ideal detection method for insulin precursors.

In this study, insulin precursors were expressed in the form of fusion proteins, and the antibodies obtained from them as antigens and the double-antis Sandwich ELISA assays were not suitable for insulin precursors produced by different production processes.

## 4. Acknowledgment

This work was financially supported by Scientific and technological research plan of Anhui Province (1604a0802101) and Major Projects of Science and Technology of Anhui Province (17030801036).

